# Forming nephrons promote nephron progenitor maintenance and branching morphogenesis via paracrine BMP4 signalling under the control of *Wnt4*

**DOI:** 10.1101/2023.11.19.567482

**Authors:** Julie L.M. Moreau, Sarah Williams, Jihane Homman-Ludiye, Andrew J. Mallett, Alexander N. Combes

## Abstract

Kidney development is known to be driven by interactions between stromal, nephron and ureteric epithelium progenitors in the nephrogenic niche. In contrast, the epithelial nephrons generated in this environment have largely been considered a product of niche rather than an active participant in the signalling interactions that maintain it. However, knockout of *Wnt4*, a gene required for nephron formation and stromal development, results in hypoplastic kidneys. We hypothesised that the forming nephron may play a role in maintaining the nephrogenic niche. In support of this hypothesis, conditional deletion of *Wnt4* from the nephron lineage resulted in nephron progenitor dispersal and death, reduced branching morphogenesis and nephron progenitor cell number. Bulk and single cell transcriptional profiling of *Wnt4* mutant kidneys revealed a downregulation of BMP signalling effectors *Id1*, *and Id3* in nephron progenitor cells, implicating *Wnt4* target BMP4 as a paracrine signal mediating feedback from the committing nephron. Recombinant BMP4 restored nephron progenitor compaction in cultured *Wnt4* mutant kidneys and blocked differentiation in wildtype controls mirroring the role of BMP7-MAPK signalling in progenitor self-renewal. Our data supports a revised model of the nephrogenic niche in which forming nephrons promote progenitor maintenance and branching morphogenesis, in part via paracrine BMP4 signalling under the control of *Wnt4*. This requirement for nephron-derived signals for maintenance of the nephrogenic niche provides new mechanistic insight into kidney morphogenesis and human renal hypodysplasia phenotypes associated with deleterious *WNT4* mutations.

## Introduction

Cycles of branching morphogenesis and nephron induction events drive kidney growth and nephron formation throughout development. Around birth, nephrogenesis ceases as the progenitor populations that comprise the nephrogenic niche differentiate. The number of nephrons that form during development varies widely in humans and mice and is an important determinant of kidney function and resistance to disease in later life. As such, the mechanisms responsible for maintaining the nephrogenic niche have been intensely studied. Nephron progenitor (NP) cells produce GDNF, BMP and FGF signals essential to maintain cell survival and branching morphogenesis in the ureteric tips that they surround. In turn, reciprocal FGF and BMP signals from the tip maintain nephron progenitor turnover (Perantoni 2005; Grieshammer et al. 2005). Regionalised WNT9B production from the base of the ureteric tip, with the aid of stromal FAT4 and DCN, triggers a mesenchymal to epithelial transition in nephron progenitor cells to form an early committing nephron (Fetting et al. 2014; Vidal et al. 2020; England et al. 2020).

Maintenance of the nephrogenic niche is known to be dependent on interactions between adjacent nephron, ureteric, and stromal progenitor cell populations (Brown et al. 2013; McMahon et al. 2008; Mugford et al. 2009). The nephrons that form in this environment are widely considered a product of the niche rather than an essential component. Yet, the early committing nephron is in close proximity to the nephron progenitor and ureteric tip populations and produces signals capable of influencing nephron progenitor fate. Indeed, complete knockout of *Wnt4*, which regulates nephron epithelialization results in kidneys with few nephrons that are also small in size. However, *Wnt4* and co-expressed signals like *Fgf8* and *Bmp4* are expressed in multiple cell types in the kidney or early embryo (Sharma et al. 2022; Mills et al. 2017; Oxburgh et al. 2005; Oxburgh et al. 2004), confounding conclusions about how their expression in the early committing nephron might affect the niche (Grieshammer et al. 2005; Perantoni 2005; Gerber et al. 2009; Mills et al. 2017; Oxburgh et al. 2005; Oxburgh et al. 2004). For example, *Wnt4* is expressed prior to development of the metanephric kidney, with lineage tracing experiments showing contributions to all major progenitor cell types (Lawlor et al. 2019; Park et al. 2012). Moreover, Wnt4 is required for proper stromal development (Itaranta et al. 2006). As such, the hypoplasia phenotype reported in *Wnt4* knockout mice may result from loss of Wnt4 action in any or all of these cell types. Mechanistically, *Wnt4* is thought to signal through a non canonical Wnt-Calcium pathway to regulate growth factors such as *Bmp4* and *Fgf8* in the early nephron, though the molecular connection between these genes and their associated phenotypes remains unclear. Finally, human mutations in *WNT4* are associated with renal hypodysplasia (Vivante et al. 2013; Zhang et al. 2021) supporting a broader role for *WNT4* in kidney development and congenital disease.

In this study we used global and conditional *Wnt4* knockout mouse models to investigate whether the early committing nephron contributes to maintenance of the nephrogenic niche. Removal of *Wnt4* from the nephron lineage impairs nephron progenitor survival, branching morphogenesis and kidney size. Transcriptional profiling, multiplexed in situ hybridisation, and ex vivo culture experiments support a mechanism where nephron progenitor maintenance is mediated in part via paracrine BMP-Smad signalling under the control of WNT4. These data support a revised model in which the early committing nephron is not simply a product of the niche but is essential to support ongoing branching morphogenesis and nephron formation.

## Results

### Nephron progenitor maintenance and branching morphogenesis are impaired in the absence of *Wnt4*

Prior analysis of *Wnt4* knockout mice identified defects in early nephron formation, stromal, and ureteric epithelium development (in UE, linage tracing in (Lawlor et al. 2019; Stark et al. 1994; Itaranta et al. 2006). Expression of *Wnt4* has been validated in each of these cell lineages. To reconcile these reports and study the role of *Wnt4* in the nephrogenic niche, we first assessed kidney development in a global *Wnt4* knockout (*Wnt4*^GCE/GCE^), where exon 2 of the *Wnt4* gene is disrupted by knock-in of a GFP-CreERT2 construct, abolishing *Wnt4* function in all cells of the embryo (Kobayashi et al. 2008). Whole mount confocal microscopy of cleared *Wnt4* knockout kidneys revealed a delay in development at embryonic day (E) 13.5 and dramatic reduction in kidney size, branching morphogenesis, and *Wnt4*-GFP expression at E14.5 and E15.5 (**Fig.1A**). Formation of early committing nephrons was severely reduced in the *Wnt4* knockout kidneys as expected (**Fig.1B**).

**Figure 1:**
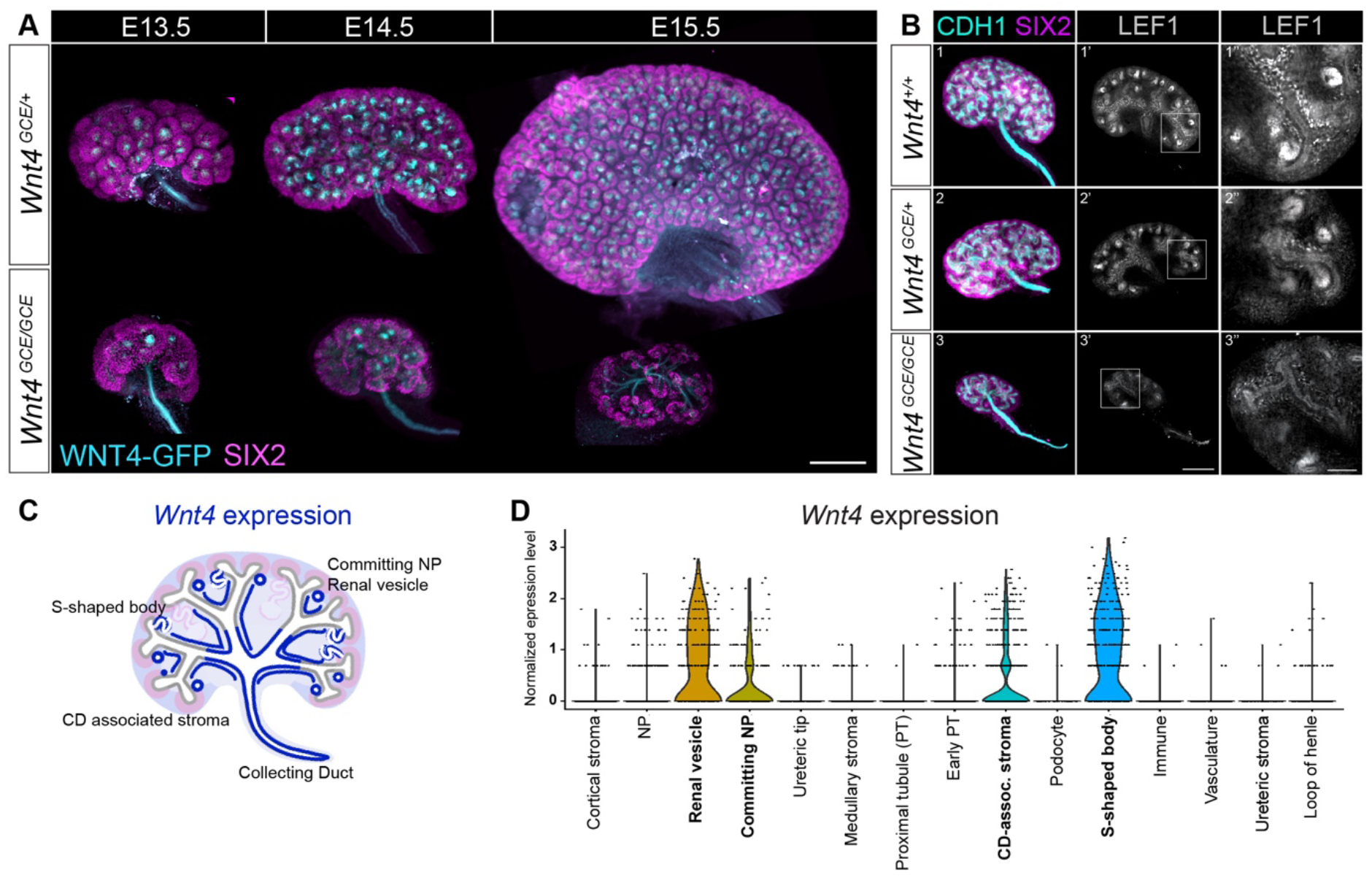
Kidney development fails in the absence of Wnt4. A) Whole organ of SIX2 and GFP antibodies stained Wnt4^GCE/+^ and Wnt4^GCE/GCE^ kidneys at 13.5 dpc, 14.5 dpc, and 15.5 dpc. Scale bar 200um. B) Whole kidneys stained with SIX2 (magenta), CDH1 (cyan) and LEF1 (gray) of Wnt 4^+/+^, Wnt4^GCE/+^ and Wnt4^GCE/GCE^ at 14.5 dpc. Scale bar 200um in 1, 1’, 2, 2’, 3 and 3’. Scale bar is 50 um in 1’’, 2’’ and 3’’. C) Illustration of Wnt4 expression domain (blue lines): Committing Nephron Progenitor, S-shaped Body, Renal Vesicle, Colleting Duct and Collecting duct associated stroma. D) Violin plot of *Wnt4* expressing population in Sc RNA-seq of E18.5 kidneys (Combes, Phipson, et al. 2019).

Although *Wnt4* is known to play an essential role in the formation of nephrons and smooth muscle cells lining the ureter (Itaranta et al. 2006), if these were the only functions of *Wnt4* then kidney morphogenesis should proceed normally until birth when impaired fluid handling and waste removal would lead to cardiovascular complications, toxicity and organ failure. Instead, the profound hypoplasia in *Wnt4*^GCE/GCE^ kidneys suggests a broader role for *Wnt4* in kidney morphogenesis. *Wnt4* is expressed in multiple cell types within the kidney renal vesicle, committing nephrons, S-shape bodies, collecting duct associated stroma, and the collecting duct itself (Fig. 1C, D). It is unclear whether the knockout phenotype is caused by loss of WNT4 function in the ureteric epithelium, nephron, stroma, or a combination thereof.

### The early committing nephron promotes kidney morphogenesis

Reciprocal signalling interactions within the nephrogenic niche play a central role in driving kidney development and the committing nephron is an integral part of the niche. As such, we hypothesised that *Wnt4* expression in the early committing nephron plays an essential and unappreciated role in maintaining the nephrogenic niche and kidney morphogenesis. To test this hypothesis, we generated a *Wnt4* conditional knockout, removing *Wnt4* function from *Six2*-expressing nephron progenitor cells (*Six2*Cre^+^, *Wnt4*^Flox/Flox^). Kidney size was reduced in *Six2*Cre^+^*Wnt4*^Flox/Flox^ embryos at E17.5 (**Fig. 2A**). The number of nephron progenitor cells and committing nephron structures per tip were also significantly reduced (**Fig.2 B**). At postnatal day (P) 5, *Six2*Cre^+^*Wnt4*^Flox/Flox^ kidneys had fewer glomeruli, disorganised proximal tubules, and were smaller than controls **(Fig.2 C)**. However, this conditional knockout phenotype was mild in comparison to the global *Wnt4* knockout. Moreover, mature nephrons were still observed to form (**Fig. 2C**), which should not be possible if *Wnt4* is effectively removed (Stark et al. 1994).

**Figure 2:**
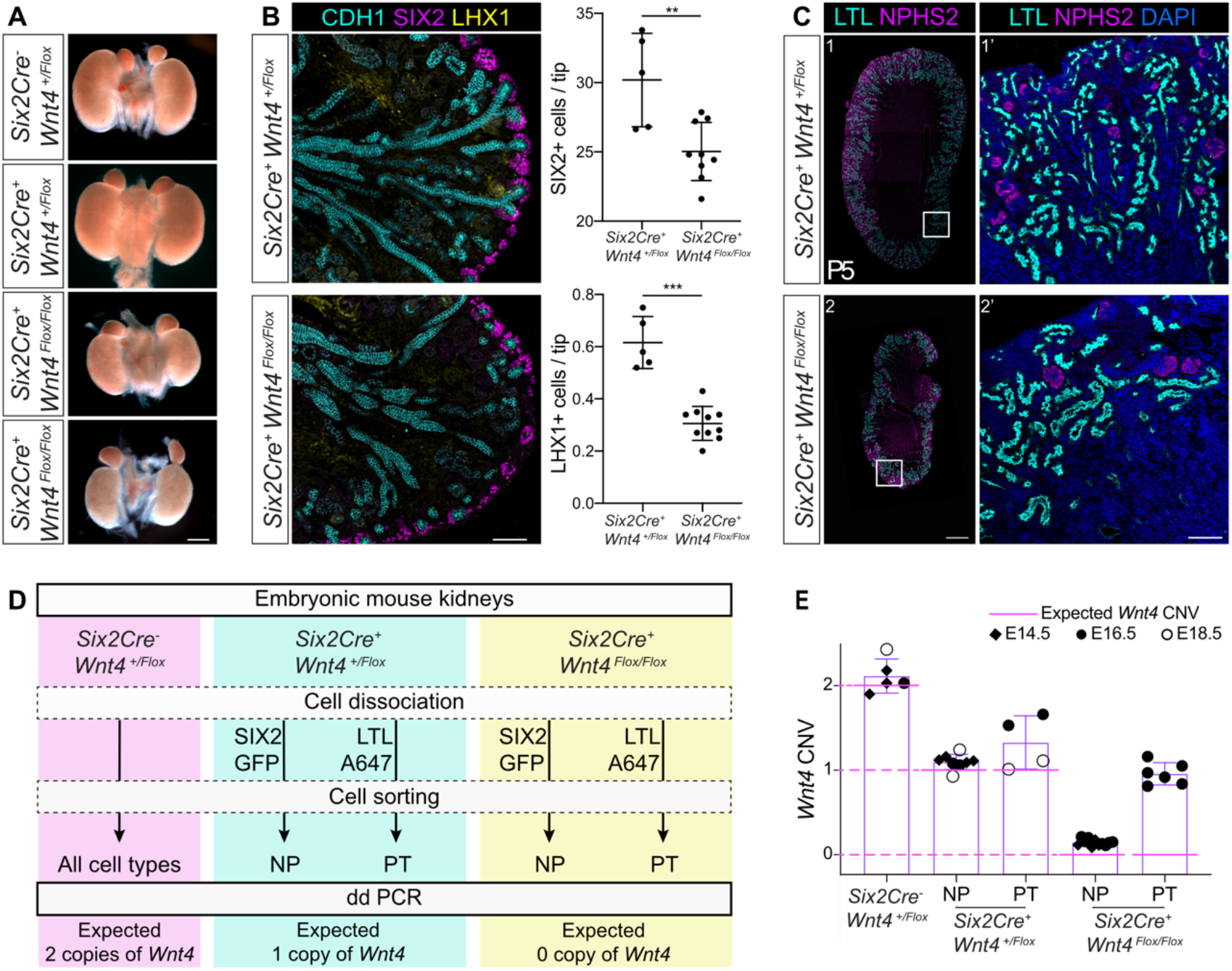
Incomplete deletion of *Wnt4* in NP derivative structures still reduces kidney size, nephron progenitor and nephron number. A) Representative brightfield pictures of kidney pairs at E17.5 of the indicated genotypes. Scale bar 500uM. B) Quantification of number of Nephron Progenitor (NP) per Ureteric Epithelium (UE) tip and LHX1 positive structures per UE tip in the Six2Cre^+^, Wnt4^Flox/+^ control and Six2Cre^+^, Wnt4 ^Flox/Flox^ conditional knock out kidneys at E17.5. Scale bar 100uM C) Representative images of a full middle section of P5 (post natal day 5) kidneys of control and Six2Cre^+^, Wnt4 ^Flox/Flox^ showing proximal tubules (LTL, Cyan, glomeruli (NPHS2, magenta) and nuclei (DAPI). Scale bar is 500uM in 1 and 2, 100uM in 1’ and 2’. D) Kidneys were isolated from controls (Six2Cre^-^, Wnt4^Flox/+^), Heterozygous (Six2Cre^+^, Wnt4^Flox/+^) and conditional Knock-outs (Six2Cre^+^,Wnt4^Flox/Flox^). Cells were dissociated and sorted for proximal tubules (PT, LTL^+^) and nephron progenitors (NP, GFP^+^). ddPCR was performed on the different sorted cell populations. E) Analysis of *Wnt4* wild type allele copy number variation (CNV) with droplet digital PCR on the sorted cell populations at E14.5, E16.5 and E18.5. The pink line represents the expected CNV for *Wnt4* WT allele presence. Six2Cre^-^ Wnt4^Flox/+^ (n=5), Six2Cre^+^Wnt4^Flox/+^ NP (n=8) and PT (n=4), Six2Cre^+^Wnt4^Flox/Flox^ NP (n=11) and PT (n=6). Graphs represent mean +/- Standard deviation.

### Incomplete deletion moderates the *Wnt4*^Flox/Flox^ phenotype

Other studies using the *Six2*Cre have reported mild or variable phenotypes, suggesting incomplete deletion of the target gene (Park, Valerius, and McMahon 2007; Chung, Deacon, and Park 2017). As such, the *Six2*Cre^+^*Wnt4*^Flox/Flox^ phenotype may be moderated by cells that retain functional copies of *Wnt4*. To test whether *Wnt4* was effectively deleted in *Six2*Cre^+^*Wnt4*^Flox/Flox^ kidneys, cells from the nephron progenitor population (GFP+) and proximal nephrons (LTL+) were isolated by fluorescence activated cell sorting from E14.5, E16.5 and E18.5 kidneys **(Fig. 2D)** and genotyped by droplet digital PCR **(Fig.2 E)**. As expected, cells isolated from control kidneys retained both copies of *Wnt4*, while only one functional copy of *Wnt4* was identified in heterozygous knockout embryos (*Six2*Cre^+^*Wnt4*^Flox/+^). In contrast, nephron progenitor cells from *Six2*Cre^+^*Wnt4*^Flox/Flox^ kidneys had a copy number variation (CNV) of 0.14 +/- 0.04, indicating retention of a functional copy of *Wnt4* in a minor fraction of nephron progenitor cells. Proximal nephron cells recovered from the same embryos retained one functional copy of *Wnt4* (CNV 0.96 +/- 0.14), suggesting that they formed prior to *Wnt4* being deleted from the progenitor pool, or from cells that retained *Wnt4*.

### The early committing nephron plays an essential and unappreciated role in maintaining the nephrogenic niche

We reasoned that breeding the conditional knockout allele (*Wnt4*^Flox/+^) onto a null allele (*Wnt4*^GCE/+^) would allow more efficient deletion of *Wnt4*. Kidneys from *Six2*Cre^+^*Wnt4*^GCE/Flox^ mice were severely hypoplastic at E17.5, with a reduced number of irregular glomeruli and tubules **(Fig.3 A)**, and an absence of tips and early committing nephrons in the nephrogenic zone compared to controls **(Fig.3 A)**. While this phenotype was more severe than the conditional knockout, the presence of nephrons and the extent of kidney development still indicates incomplete deletion of *Wnt4*. Nonetheless, these results demonstrate that expression of *Wnt4* in the early committing nephron is required to maintain the nephrogenic niche.

**Figure 3:**
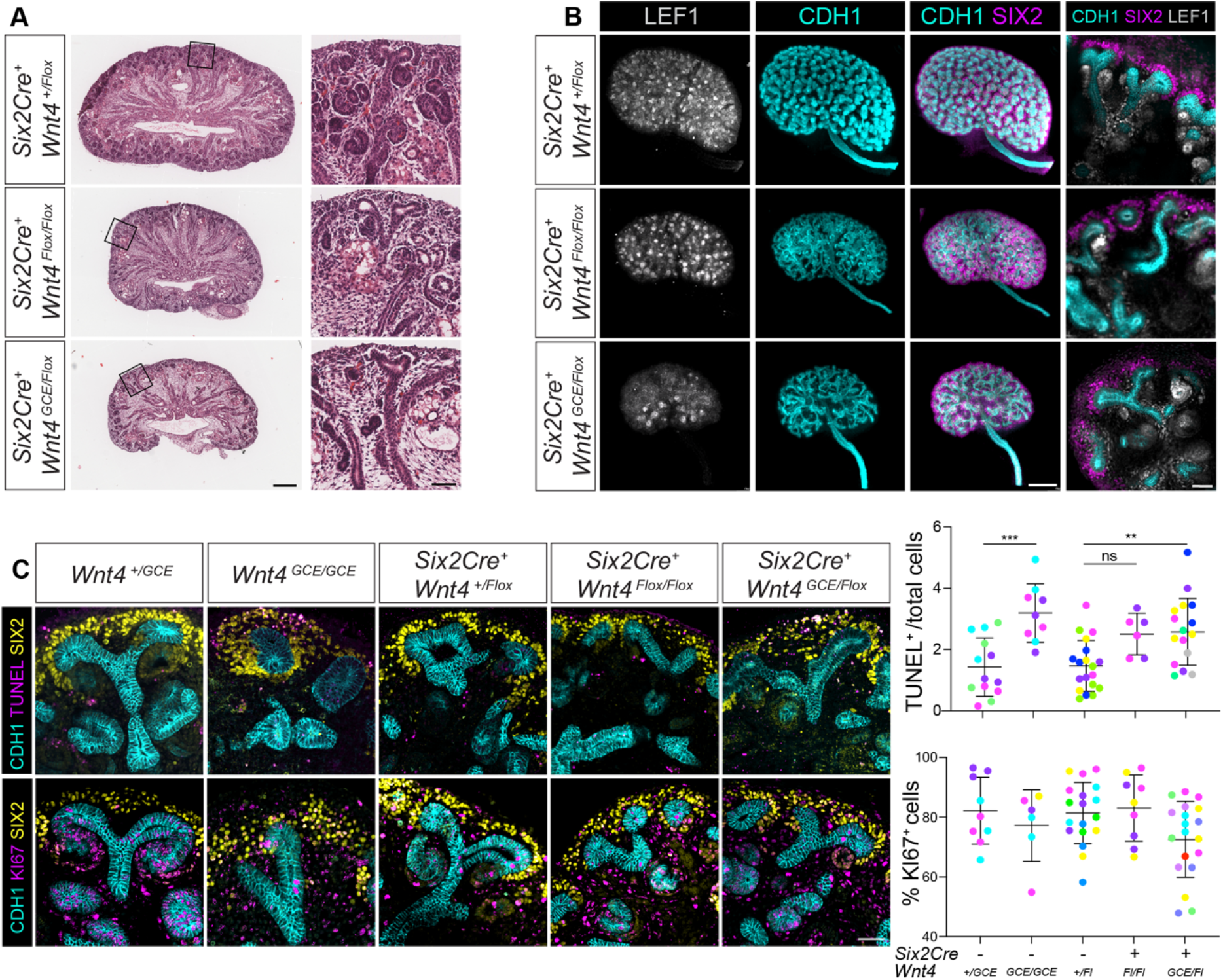
Conditional knockout of *Wnt4* in NP reduces kidney size, nephron number and nephron progenitor via apoptosis. A) Representative images of H&E stained middle section of E17.5 kidneys from Six2Cre^+^,Wnt4^Flox/+^ control, Six2Cre^+^,Wnt4^Flox/Flox^ and Six2Cre^+^,Wnt4^GCE/Flox^ conditional KOs. Scale bar 200 um and close up of niches 50 um. B) Representative images of whole mount kidneys stained with LEF1 (grey), ECAD (Cyan) and SIX2 (Magenta) at E14.5 of the indicated genotypes Scale bar 200 um. Close up of niches with merged channels is also represented for all the listed genotypes. Scale Bar 200uM for the whole kidneys and 50uM for the niches close ups.C) Quantification of apoptosis and proliferation using immunofluorescence in nephrogenic niches is the listed genotypes. TUNEL positive cells where quantified on each picture and normalised to the total number of cells using DAPI (staining not represented). The same methodology was applied to KI67 positive cells and represented as a percentage. Each dot represents a quantified picture, each colour represents a sample. One way ANOVA and Tukey’s multiple comparison test were performed on samples. ns: non significant, **P<0.01, *** P<0.001. Scale bar 50uM.

Kidneys from *Six2*Cre^+^*Wnt4*^Flox/Flox^ and *Six2*Cre^+^*Wnt4*^GCE/Flox^ mice were assessed at E14.5 to determine the cellular mechanisms underlying the hypoplastic phenotype. Variability was noted in both conditional knockouts, which we expect to arise from incomplete deletion of the floxed *Wnt4* allele/s (Fig.3 B). Despite this, branching was impaired, the number of committing nephrons were reduced and a decrease of nephron progenitors was also evident **(Fig.3 B)**. Cell death was quantified in the nephrogenic zone from the global and conditional knockouts using the TUNEL assay **(Fig.3 C)** (n=3-7 per condition). Cell death was significantly increased in *Wnt4*^GCE/GCE^ global knockout embryos (*Wnt4*^GCE/GCE^ n=3 3.19 +/- 0.95 and *Wnt4*^GCE/+^ n=4 1.43 +/- 0.98) and in the *Six2*Cre^+^*Wnt4*^GCE/Flox^ conditional knockout (*Six2*Cre^+^*Wnt4*^+/Flox^ n=6 1.47 +/- 0.83, *Six2*Cre^+^*Wnt4*^Flox/Flox^ n=6 2.5 +/- 0.67 and *Six2*Cre^+^*Wnt4*^GCE/Flox^ n=5 2.57 +/- 1.09) **(Fig.3 C)**. Despite the dramatic difference in kidney size at later stages, the proportion of cells expressing proliferation marker KI67 was not significantly different between genotypes at E14.5 but trended down in *Wnt4*^GCE/GCE^ and *Six2*Cre*Wnt4*^GCE/Flox^ **(Fig.3 C)**.

### Bulk RNAseq identifies increased stromal signatures in *Wnt4* knockouts, confirms altered branching and nephrogenesis

Knockout of *Wnt4* from the nephron lineage demonstrates a new role for *Wnt4* or other downstream signals in the early committing nephron in maintaining the nephrogenic niche. To examine the transcriptional response to loss of *Wnt4*, bulk RNA sequencing (RNAseq) was performed on *Wnt4*^GCE/GCE^ kidneys at E14.5 and *Six2*Cre^+^*Wnt4*^Flox/Flox^ kidneys at E14.5, E17.5 (Supplementary Table 1).

289 genes were differentially expressed between *Wnt4*^GCE/+^ and *Wnt4*^GCE/GCE^, of which 101 were down regulated in *Wnt4*^GCE/GCE^ and 188 were upregulated (FDR cut-off 0.05 and abs logFC 0.585/1.5x). Differentially expressed gene lists were cross-referenced to single cell RNAseq data from wildtype E18 kidney to identify changes associated with specific cell types. As expected, 84% of downregulated genes were expressed in structures that derive from the early committing nephron, which are largely absent in knockout embryos. Downregulated genes included *Wnt4*, and genes proposed as *Wnt4* targets such as *Pax8*, *Lhx1,* and *Fgf8* **(Fig. 4A)** (Valerius and McMahon 2008) as well as nephron progenitor cell markers *Cited1* and *Itga8*. Surprisingly, 81% of the upregulated genes were associated with stromal cell populations (153/188) including *Foxd1*, *Pdgfra*, *Actc1* and *Crabp1* **(fig. 4A)**.

**Figure 4:**
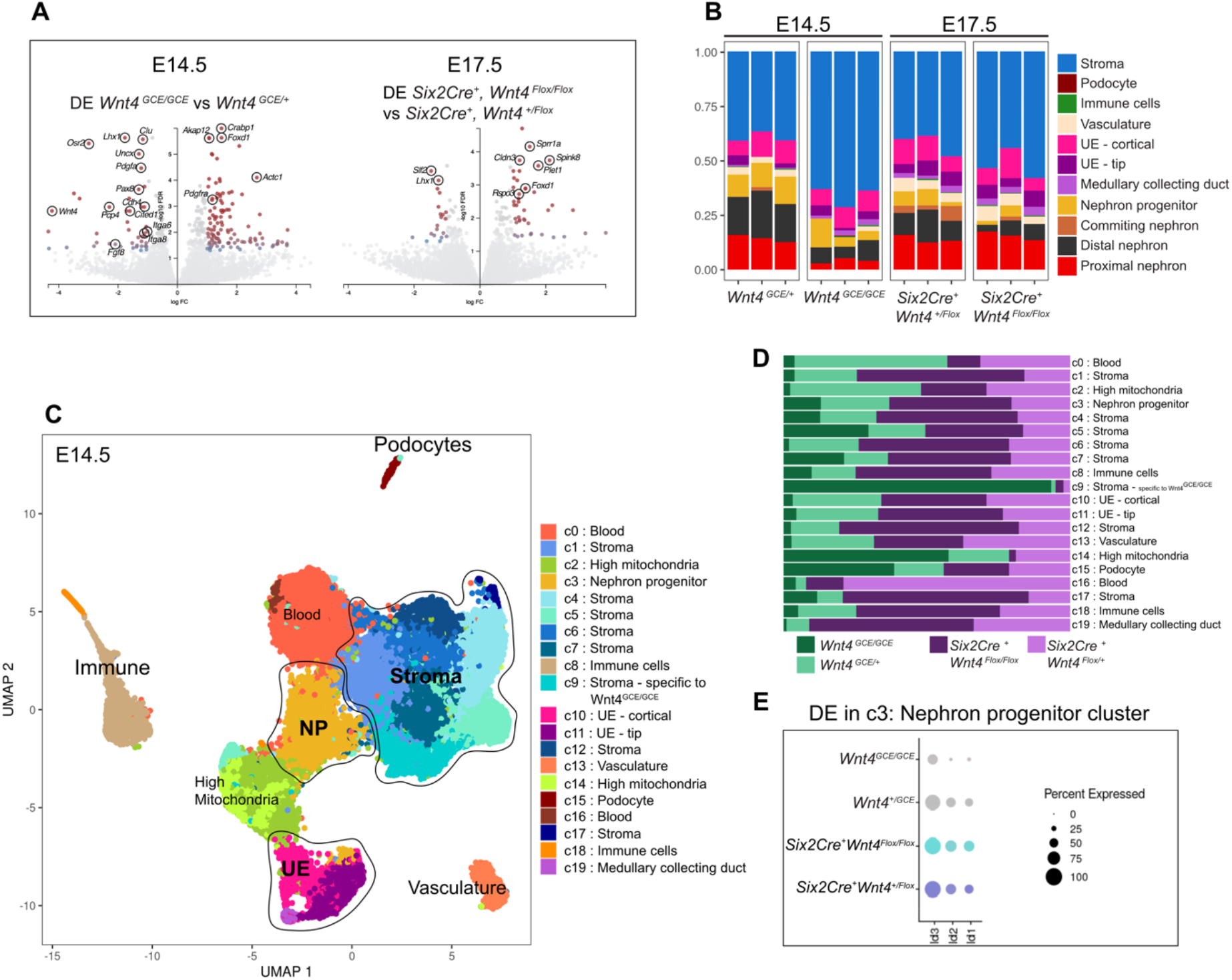
Six2-derived structures and stromal cell populations are transcriptionally affected in absence of *Wnt4*. A) Differential expression volcano plots of Bulk RNA sequencing performed on Wnt4^GCE/+^, Wnt4^GCE/GCE^ at E14.5 displaying the 101 genes downregulated (red dots, left side of the plot), and 188 upregulated (red dots, right side of the plot). Similar experiment and presentation shown on the second volcano plot of differentially expressed genes between Six2Cre^+^, Wnt4^Flox/+^ and Six2Cre^+^, Wnt4^Flox/Flox^ at E17.5. n=3 per genotype. B) Bar graph showing the 3 separate samples of each genotype using deconvolution of the Bulk RNA sequencing results with E18.5 Single Cell RNA sequencing datasets previously published in (Combes et al., 2019). Each colour represents a cluster. C) UMAP of Single Cell RNA sequencing performed on Wnt4^GCE/+^, Wnt4^GCE/GCE^, Six2Cre^+^ Wnt4^Flox/+^ and Six2Cre^+^ Wnt4^Flox/Flox^ kidneys at E14.5. Total cell number: 19360. D) Horizontal bar graph representing the proportion of cells of each genotype per cluster from the single cell RNAseq. E) Dot plot representing three DNA-binding protein inhibitor Id1, Id2 and Id3 genes that are differentially expressed in the nephron progenitor c3 cluster. These 3 markers are downregulated in Wnt4^GCE/GCE^ and Six2Cre^+^, Wnt4^Flox/Flox^ compared to their relevant controls at E14.5. See Fig S1 and supp tables 1, 2, 3 and 4 for full lists)

**Figure 5:**
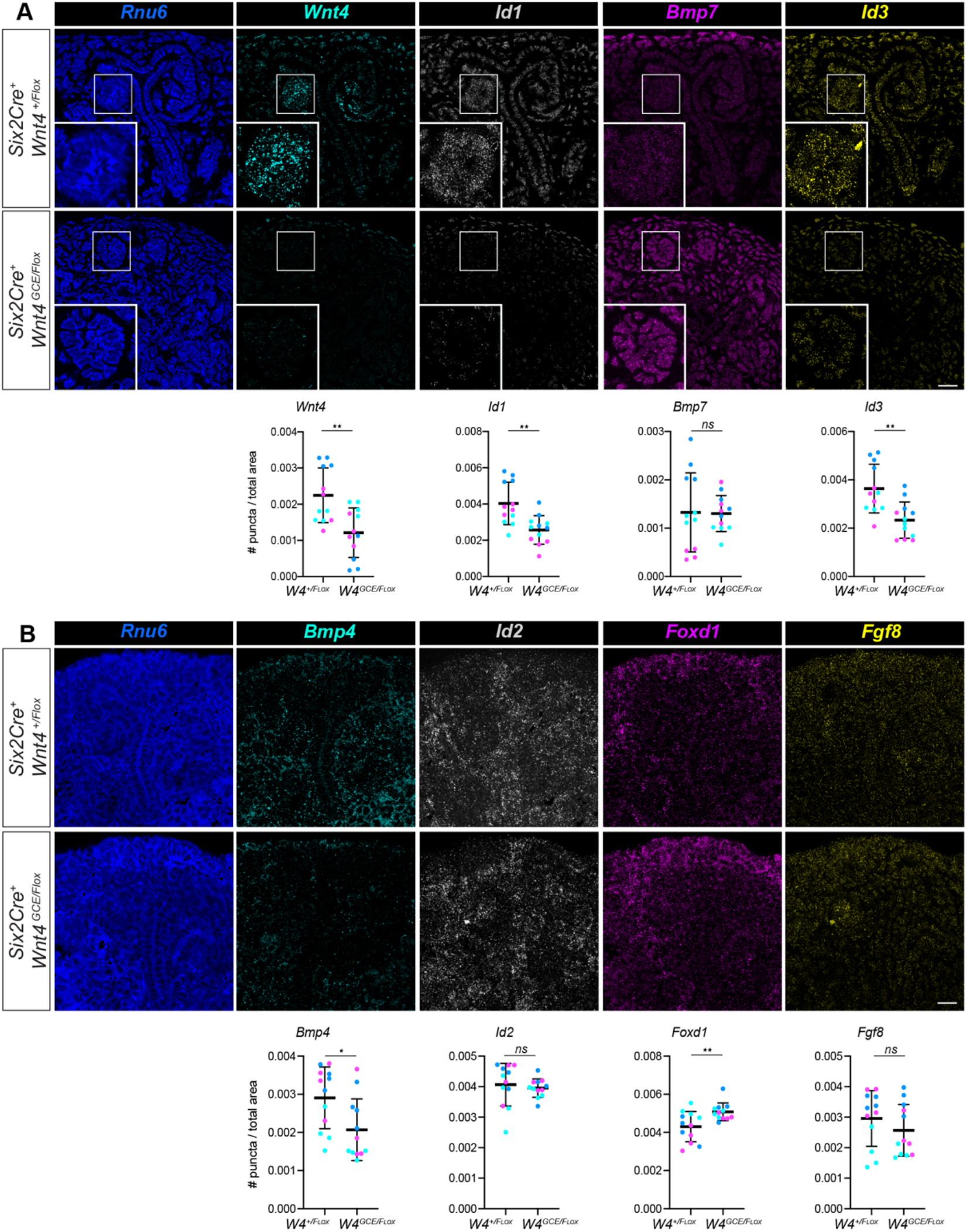
Conditional knockout of Wnt4 with in NP decreases Id1, id2, Bmp7 and increases Foxd1 in nephrogenic niches. A) Quantification of mRNAs using Hybridisation chain reaction (HCR) of the listed RNAs relative to the reference RNA, *Rnu6* in nephrogenic niches. The images are representative of each staining. The zoom represents each staining in the committing nephron. B) Second set of experiment involving quantification of mRNAs using Hybridisation chain reaction (HCR) of the listed RNAs relative to the reference RNA, *Rnu6*. The images are representative of each staining. Each dot represents a quantified picture, each colour represents a sample. Student T-test were performed on samples. ns: non significant, *P<0.05, **P<0.01, *** P<0.001. Scale bar 20uM.

**Table 1:** Antibodies used for immunodetection.

| Target | Catalogue number | Species | Supplier | Dilution |
| --- | --- | --- | --- | --- |
| **PRIMARY ANTIBODIES** |  |  |  |  |
| SIX2 | 11562-1-AP | Rabbit | Proteintech | 1:300 |
| LEF1 | 2230 | Rabbit | Cell Signalling | 1:300 |
| CDH1 | AF748 | Goat | R&D | 1:500 |
| NPHS2 | P0372 | Rabbit | Sigma Aldrich | 1:300 |
| LTL | B-1325 | N/A Biotin | Vector Laboratories | 1:600 |
| LHX1 | 4F2 | Mouse | DSHB | 1:20 |
| KI-67 | 13-5698-82 | N/A Biotin | Invitrogen | 1:200 |
| GFP | ab13970 | Chicken | Abcam | 1:100 |
| **SECONDARY ANTIBODIES** |  |  |  |  |
| Donkey anti-Rabbit Alexa Fluor™ 568 | A10042 |  | Invitrogen | 1:1,000 |
| Donkey anti-Rabbit Alexa Fluor™ Plus 488 | A32790 |  | Invitrogen | 1:1,000 |
| Donkey anti-Rabbit Alexa Fluor™ 568 | A10042 |  | Invitrogen | 1:1,000 |
| Donkey anti-mouse Alexa Fluor™ Plus 647 | A32787 |  | Invitrogen | 1:1,000 |
| Donkey anti-Goat Alexa Fluor™ Plus 488 | A32814 |  | Invitrogen | 1:1,000 |
| Streptavidin, Alexa Fluor™ 647 | S21374 |  | Invitrogen | 1:2,000 |
| Streptavidin, Alexa Fluor™ 488 | S11223 |  | Invitrogen | 1:2,000 |
| Streptavidin, Alexa Fluor™ 568 | S11226 |  | Invitrogen | 1:2,000 |
| **NUCLEAR STAIN** |  |  |  |  |
| DAPI | 10236276001 |  | Sigma-Aldrich | 1:1,000 |

**Table 2:**
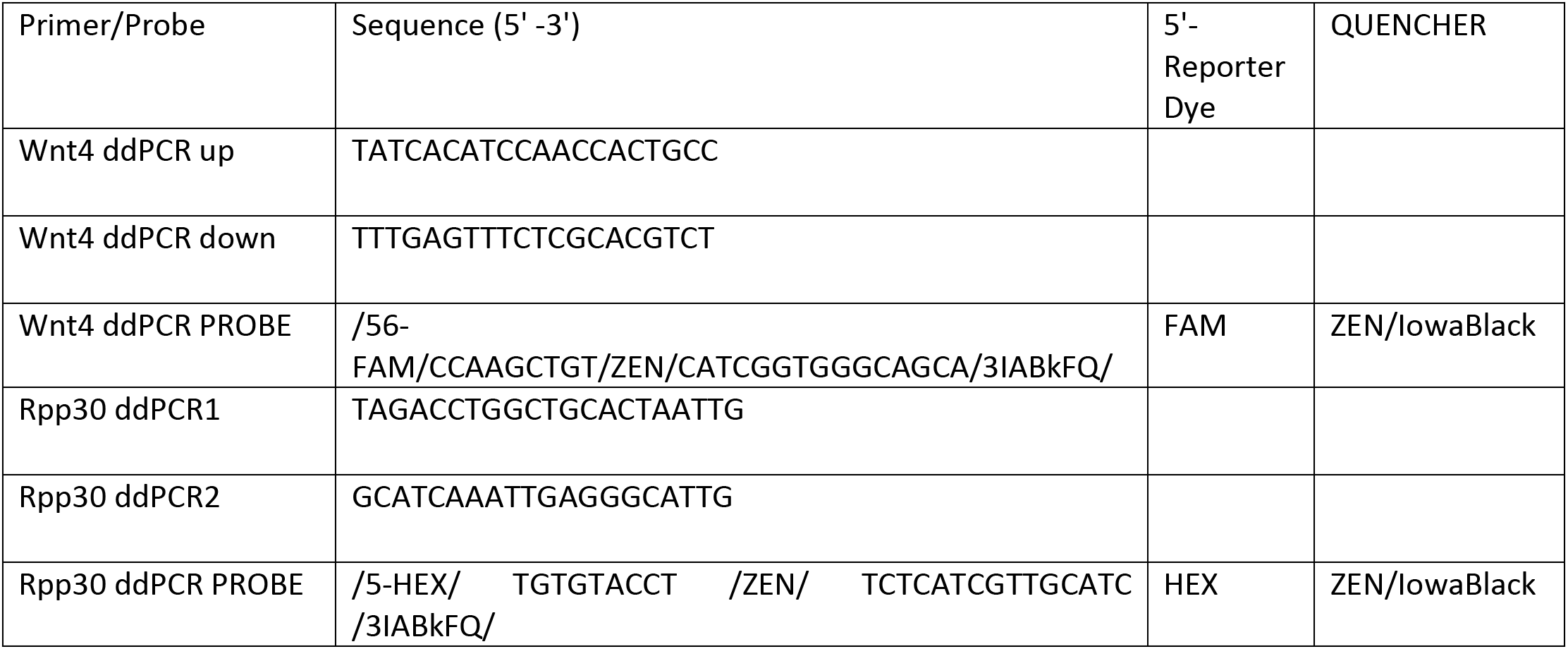
Primers and probes used for Wnt4 copy number variation assay by ddPCR.

No transcriptional differences were found between *Six2Cre^+^Wnt4^Flox/Flox^*and *Six2Cre^+^Wnt4^Flox/+^* at E14.5. However, 206 genes were differentially expressed at E17.5, reflecting the increased deletion and severity of the phenotype with time (Table S1). Downregulated genes were again associated with reduced nephron formation, with ∼5% associated with stromal cell populations **(Fig. 4A)**. Consistent with the *Wnt4*^GCE/GCE^ data, a third of upregulated genes in *Six2Cre^+^Wnt4^Flox/Flox^* were stromal cell markers such as *Foxd1* and surprisingly, 56% were associated with cortical and medullary collecting duct cell types such as *Spink8*, *Krt18*, *Fxyd3* or *Krt19* **(Table S1 and Fig. 4A)**. The increase in medullary collecting duct markers may indicate a compromised ureteric tip progenitor cells state, and enhanced differentiation of cortical and medullary collecting duct, consistent with the impaired branching phenotype observed in these embryonic kidneys.

Overall, these findings suggest that *Wnt4* or other signals in the early committing nephron influence the maintenance and differentiation of progenitor cells in the stroma and ureteric epithelium.

### Wnt4 knockout scRNAseq identifies downregulation of *Id1, Id2, Id3* in nephron progenitor cells

We next performed single cell RNA sequencing (scRNAseq) on *Wnt4* heterozygous and knockout kidneys at E14.5 to assess how individual cell types within the nephrogenic niche respond to loss of *Wnt4* from the early committing nephron. Three kidneys of each genotype were dissociated, pooled and sequenced on the 10x Chromium platform. Quality control measures included doublet removal, cell filtering by UMI counts and normalisation for cell cycle stage. Unsupervised clustering of the remaining 19360 cells resulted in 19 distinct populations, which were assigned identities based on marker analysis **(Fig 4C, Supp Fig.1 and Supplementary Table 2)** and via label transfer from our reference dataset (Combes, Phipson, et al. 2019). Clusters representing nephron progenitor, ureteric tip, and cortical stroma were identified, in addition to vascular, blood and immune cell populations **(fig. 4C)**. One cluster with a high mitochondrial gene content was also recovered, which was primarily composed of cells from *Wnt4* knockout kidneys **(c2 and c14, fig. 4D and table S2)**. Given that multiple *Wnt4^GCE/GCE^* and control samples were collected and processed in parallel for this experiment, this may reflect a cell stress phenotype in the knockout rather than a dissociation artefact. Cellular stress is consistent with the elevated levels of apoptosis seen in the nephrogenic niche.

Differential expression analysis of cell types between control and Wnt4 KO kidneys identified a consistent increase in mitochondrial gene expression in the *Wnt4* KO with elevated expression of genes such as *mt-Atp6*, mt-Co2 or *mt-Co3*. Beyond this recurrent signature, elevated labels of *Foxd1* were observed in Wnt4KO stromal progenitors, while ureteric tip markers did not show significant differences **(c11, Table S5)**. However, genes best known as BMP-SMAD signalling effector genes *Id1*, *Id2*, and *Id3* were significantly(?) downregulated in the nephron progenitor cell cluster in both Wnt4^GCE/GCE^ and *Six2*CreWnt4^Flox/Flox^ where they were decreased by 25% (0.75 logFC, adj p = X) **(Fig 4E, Table S5)**. Genes previously identified as *Wnt4* target genes like *Fgf8* were not identified as differentially expressed. While this could be due to a lack of sensitivity in the scRNAseq experiment or failure to pass the statistical thresholds for differential expression, changes in *Id1, Id2, Id3* and stromal progenitor genes like *Foxd1* are robust outcomes that were identified in bulk and single cell experiments.

### Forming nephrons extend the lifespan and output of the nephrogenic niche via BMP signalling

*Wnt4* is well established as a regulator of nephron epithelialization, with prior analysis suggesting that *Wnt4* regulates *Bmp4* and *Fgf8* expression in the early committing nephron (Torban et al. 2006; Kim et al. 2006; Grieshammer et al. 2005). Furthermore, *Bmp7*, also part of the TGFB signalling pathway is strongly expressed in metanephric mesenchyme and all the derivative structures (Muthukrishnan et al. 2015; Motamedi et al. 2014; Blank et al. 2009; Oxburgh et al. 2005; Godin et al. 1998). While *Bmp7*, *Bmp4* and *Fgf8* were not identified as differentially expressed in our single cell analysis, transcriptional effectors of BMP-SMAD signalling *Id1*, *Id2*, and *Id3* were downregulated in nephron progenitor cells. BMP-Smad signalling is essential for maintenance of the nephrogenic niche, with ligands BMP7 expressed in nephron progenitor cells and the ureteric tip (Oxburgh et al. 2005), and BMP4 expressed in the early committing nephron and derivatives (Oxburgh et al. 2014). Loss of Bmp ligands or SMAD effectors from the entire kidney or stromal and nephron progenitor cells triggers apoptosis in the nephrogenic niche. As such, *Wnt4* may regulate nephron progenitor survival and promote ureteric branching through BMP4. To test this hypothesis and for stromal changes, we assessed expression of *Wnt4*, *Bmp4*, *Bmp7*, *Id1*, *Id2*, *Id3,* and *Foxd1* utilising a single molecule *insitu* hybridisation method, hybridisation chain reaction (HCR). Paraffin sections of wild type and *Six2CreWnt4^GCE/Flox^*at E17.5 (n=3 each) were hybridised with probes against *Wnt4*, *Id1*, *Bmp7*, *Bmp4*, *Foxd1*, *Id1*, *Id2*, and *Id3* and quantified in the niche using a house keeping gene*, Rnu6. Wnt4* was significantly reduced in the committing nephron of knockout and conditional mutatnts compared to controls, validating the experimental approach. *Foxd1* was significantly upregulated in the conditional mutant compared to the wild type (n=3 for each, P= 0.008), confirming our earlier findings. Next, we assessed *Id1, Id2, Id3*, finding significant reductions in *Id1* (n=3 for each, P= 0.0016) and *Id3* (n=3 for each, P= 0.0016) recorded in *Six2CreWnt4^GCE/Flox^*, while *Id2* was not significantly affected in this experiment (n=3 for each, P= 0.62). Surprisingly, *Bmp7* and *Fgf8* were not significantly changed even though the committing nephrons structures are severely affected. Finally, *Bmp4* was downregulated in *Six2CreWnt4^GCE/Flox^* (n=3 for each, P= 0.019) supporting a role for the early committing nephron in activating TGFB signalling in nephron progenitor cells to maintain the nephrogenic niche.

### BMP4 mediates NP compaction in the absence of WNT4

BMPs have established roles in nephron progenitor self-renewal (BMP7-MAPK), differentiation (BMP7-SMAD)(Brown et al. 2013; Oxburgh et al. 2005). Too little BMP signalling compromises nephron progenitor cell survival as knockout of BMP4, BMP7 or their receptors is associated with loss of nephron progenitors, while excess BMP signalling can also trigger apoptosis (Motamedi et al. 2014; Michos et al. 2004). Our findings implicate BMP4 as a paracrine signal to regulate nephron progenitor survival. To test if BMP4 recues the nephron progenitor phenotype, we cultured E12.5 WT and Wnt4^GCE/GCE^ kidneys for 48h in presence of recombinant BMP4 at 100ng/ml as previously published(Mills et al. 2017) and stained for SIX2, ECAD and LHX1 **(Fig.6)**. In vehicle controls, nephron progenitor cells were clustered around the tips of wildtype kidneys but dispersed from the tips in *Wnt4* knockouts (mean 26μm control vs 47μm *Wnt4^GCE/GCE^*). Addition of BMP4 into the media restored nephron progenitor compaction around the tips in *Wnt4* knockouts (mean 47μm to 25μm with BMP4). As BMP4 was added to the entire culture it is not clear if the rescue of NP compaction was a direct effect of BMP4 on nephron progenitor cells or if it is an indirect effect via a change of in the ureteric tip. Recombinant BMP4 has been reported to promote differentiation of early collecting ducts into ureter-like epithelial tubules (Mills et al. 2017). This phenomenon could promote loss of tip identity and result in a failure to maintain nephron progenitor cells and may explain why some tips in knockout and control kidneys supplemented with BMP4 lost all surrounding nephron progenitor cells. Interestingly BMP4 also impaired nephron formation in wildtype controls, mirroring the effect of BMP7-MAPK signalling to maintain self-renewal and block differentiation. These experiments support a role for paracrine BMP4 signalling in regulating nephron progenitor compaction and self-renewal.

**Figure 6:**
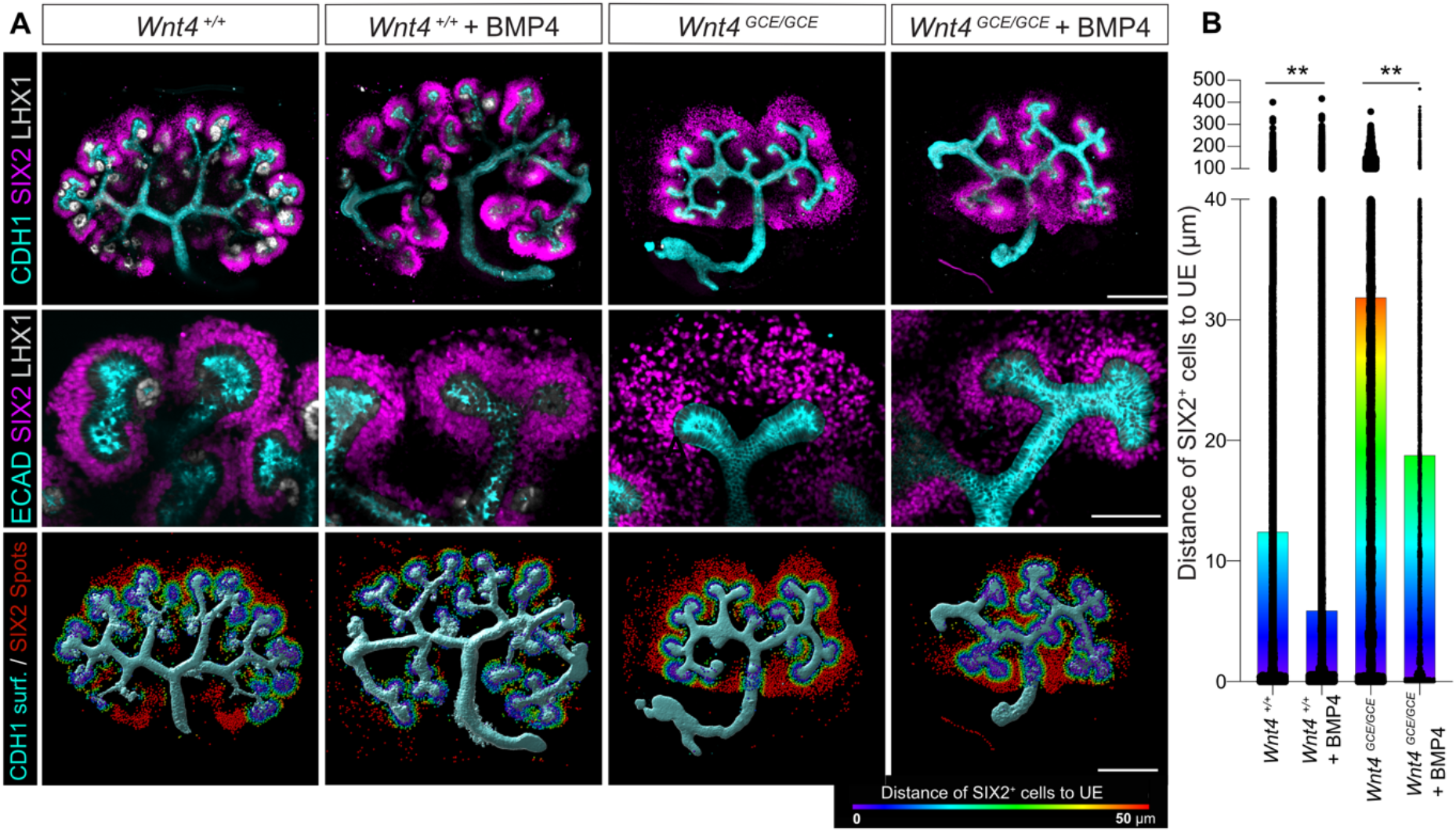
BMP4 rescues the compaction of nephron progenitor around the ureteric epithelium in absence of *Wnt4*. A) Representative images of E12.5 kidneys grown on transwell for 48h for the listed genotypes in the listed conditions. BMP4 ligand was added at 100 ng/ml. Kidneys were stained for nephron progenitors (SIX2, Magenta), ureteric epithelium (ECAD, Cyan) and early committing nephrons (LHX1, white). Scale bar 400um in whole kidney maximum intensity projection and 100um on zoomed in of nephrogenic niches. B) Quantification graph representing the distance of NP to UE using Imaris. Wnt4^+/+^ and Wnt4^+/+^ + BMP4 (n=5 for both) and about 30 000 cells quantified respectively. Wnt4^GCE/GCE^ (n=3, about 18 000 cells) and Wnt4^GCE/GCE^ + BMP4 (n=4, about 24 000 cells quantified) The rendered SIX2 spots are colour coded according to the distance to UE from 0 to 50uM. ** P<0.01

## Discussion

The developing mouse kidney is a classic model of inductive interactions in vertebrate organogenesis (Grobstein 1955). Early microdissection and co-culture studies inferred a requirement for mesenchymal signals to induce outgrowth of the ureteric epithelium, and a corresponding requirement for ureteric signals to induce nephron formation (Grobstein Inductive interaction. 1955). In time major signals responsible for maintaining branching morphogenesis and nephron induction were elucidated and models of kidney development assumed nephron progenitors were solely dependent on signals from the ureteric epithelium until signals produced by stromal progenitors were found to promote nephron progenitor differentiation, forcing a revision of this model (Fetting et al. 2014; Carroll and Das 2013). In this study we demonstrate that the early committing nephron plays an essential role in maintaining nephron progenitor survival and proliferation, in part through BMP4 signalling downstream of Wnt4.

Previous work in the field has focussed on nephron progenitor - ureteric tip interactions as necessary for maintaining branching morphogenesis with recent support for stromal progenitors providing additional feedback by regulating nephron progenitors differentiation. If true, then kidney development should progress normally in the absence of nephrons until birth when nephron function is required for life. However, our data shows that kidneys lacking *Wnt4* from the nephron lineage alone result in a hypoplastic phenotype with impaired nephron progenitor maintenance and reduced branching morphogenesis. Thus, *Wnt4* or other signals from the early committing nephron are required to maintain the nephrogenic niche.

Although hypoplastic phenotypes were previously noted to result from global *Wnt4* deletion, the simultaneous loss of this gene from sites of expression in the early embryo, stroma and ureteric epithelium clouded interpretation of this phenotype. Here we demonstrate that loss of *Wnt4* from the nephron lineage results in hypoplasia. Similarly, while other potential targets of *Wnt4* in the early committing nephron including *Fgf8* and *Bmp4/7/SMADs* are associated with hypoplastic phenotypes, single cell gene expression profiling studies have revealed that these ligands are also expressed in other cell types. For example, FGF8 is expressed in NP as well as PTA, Bmp7 in NP and UT, only BMP4 is exclusively expressed in RV.

Our data suggests new feedback between the committing nephron and the cortical stromal progenitor population marked by *Foxd1*. It is still unclear whether the forming nephron suppresses stromal progenitor identity, whether forming nephrons promote stromal differentiation, or whether loss of nephron progenitors allows expansion of the cortical stroma. Indeed, it has been shown that, in absence of FAT4, a Cadherin present in cortical stroma, the size of NP pool around the UE tip is increased (Bagherie-Lachidan et al. 2015). In our bulk RNA sequencing, we see an increase of *Fat4* in Six2Cre^+^ Wnt4^Flox/Flox^, suggesting an explanation why the Nephron progenitor pool loss in the niche. Furthermore, loss of the stromal transcription factor gene Foxd1 results in reduced branching and an expanded Cap mesenchyme (Hatini et al., 1996). Overall, the phenotype we observe in absence of *Wnt4* or committing nephron on UE branching, NP maintenance could be amplified by a direct or indirect effect via the cortical stroma. However, most reported stromal expansion phenotypes are reported to enhance nephron progenitor differentiation, which may have unintended consequences in the *Wnt4* knockout where nephron differentiation fails to progress. *Fgf8* is regulated by *Wnt4* and expressed at the highest levels in pre-tubular aggregates and early renal vesicles (Perantoni 2005; Combes, Phipson, et al. 2019; Grieshammer et al. 2005). However, recent studies have identified *Fgf8* expression in nephron progenitor cells, where *Wnt4* is not expressed at high levels. *Fgf8* is required to regulate the survival of nephron progenitor cells (Grieshammer et al. 2005). Similarly, *Fgf9* and *Fgf20* are expressed in NPs and have established roles in survival and self-renewal. Another recent study highlighted the significant chemokinetic effect of FGF8 and its importance in the condensation of nephron progenitor cells around the ureteric tip (Sharma et al. 2022). Specifically, when *Fgf8* was deleted in *Wnt4* positive cells, it resulted in smaller kidneys, fewer nephrons, and disorganized nephron progenitors, which is consistent with the findings of the present study in the absence of *Wnt4* in the niche. We did not identify significant changes in *Fgf8* expression by scRNAseq or HCR but it’s likely that *Fgf8* plays a role in the *Wnt4* knockout phenotype.

Our scRNAseq experiments identified changes in *Id1, Id2,* and *Id3* expression in nephron progenitor cells. These genes are best known as transcriptional effectors of BMP-SMAD signalling, which has shown to regulate nephron progenitor differentiation under the control of BMP7 (Lindstrom et al. 2015; Oxburgh et al. 2011; Blank et al. 2008; Oxburgh et al. 2005; Oxburgh et al. 2004). As such, we investigated whether proposed *Wnt4* target *Bmp4* may be acting as a paracrine signal. BMP4 is a member of the TGF-β superfamily and is known to play diverse roles in embryonic development. In the nephrogenic zone, BMP4 is expressed in the renal vesicle (Oxburgh et al. 2011) and acts as a paracrine factor, influencing adjacent nephron progenitors through direct cell-cell interactions. *Bmp4* mRNA was reduced by HCR and recombinant BMP4 both restored NP compaction in *Wnt4* mutants and blocked NP differentiation in wildtype controls - mimicking the effect of BMP7-MAPK. While this is apparently at odds with the link between *Id* genes and the pro-differentiation BMP-SMAD pathway, *Id* genes have also been characterised as downstream targets of other signalling cascades including the MAPK pathway(Lim and Wu 2005; Roschger and Cabrele 2017). While further studies will be required to determine which pathways are disrupted by the absence of *Wnt4*, our results are consistent with an essential role for paracrine BMP4 signalling to maintain nephron progenitor cell survival. We propose that paracrine BMP4 acts in concert with BMP7 to promote self renewal and maintain a healthy progenitor cell population. While we cannot rule out a role for paracrine BMP4 to directly affect the ureteric tip, the absence of significantly downregulated tip genes in the bulk and single cell experiments suggest that impaired branching morphogenesis is caused by the loss of nephron progenitor cells in this model.

Finally, our study suggests that, in absence of WNT4, Nephron progenitors present decreased *Id1* and *Id3* transcripts. Id (inhibitor of DNA binding proteins) genes are known to be regulating tissue specific cell proliferation, differentiation, apoptosis, and fibrotic processes (Jen, Manova, and Benezra 1996). These two genes are also known to be direct targets of BMP4 signalling in human trabecular meshwork cells and embryonic stem cells (Mody, Wordinger, and Clark 2017; Hollnagel et al. 1999). In the developing kidney, *Id1* and *Id3* are expressed in the nephrogenic zone (Allen Brain atlas and(Combes, Zappia, et al. 2019)) and are likely to be activated through BMP4, SMAD1/5/8 signalling pathway as it does in proximal tubular cells after kidney injury (Vigolo et al. 2019). The *Id* genes are likely to be involved in nephron progenitor maintenance and differentiation that is altered in absence of committing nephrons.

Our study also raises the question of whether the PTAs population plays a role in contributing to the development of multiple nephrons. Our analysis of conditional *Wnt4* knockout using the *Six2TGC-Cre* allele revealed incomplete deletion of *Wnt4* and a less pronounced phenotype. This suggests the possibility of positive selection and clonal expansion of the rare *Wnt4*-expressing cells, which could contribute to the neighbouring PTAs. These open questions will need to be addressed in future studies.

While the hypoplastic phenotypes of global *Wnt4, Fgf8* and *Bmp4* knockout mice have hinted at a broader role for the early committing nephron in kidney development, this possibility has been confounded by multiple sites of expression and pleiotropic phenotypes. As such, countless reviews declare that kidney development is driven by reciprocal interactions between nephron progenitor cells, the ureteric tip, and the surrounding stroma. Our study demonstrates that the early committing nephron is required to maintain the nephrogenic niche, in part via paracrine BMP4 signalling. This finding provides a mechanism for human renal hypoplasia phenotypes associated with deleterious *WNT4* mutations (Zhang et al. 2021)and adds another reciprocal signalling partner to the nephrogenic niche.

## Methods

### Mouse lines

All animal experiments were assessed and approved by the Monash Animal Ethics Committee MARP-2 (22165 and 22271) and were conducted in accordance with applicable Australian laws governing the care and use of animals for scientific purposes.

Wnt4 knock out embryos were produce by mating two Wnt4^GCE/+^ (B6.Cg Wnt4^tm2(EGFP/cre/ERT2)Amc^/J, (IMSR_JAX:032489) (Kobayashi et al. 2008). Conditional knock out in nephron progenitor lineage were produced by mating Six2^TGC^, Wnt4^Flox/+^ (Six2-TGC^tg^, IMSR_JAX:009606) (Kobayashi et al. 2008) with Wnt4^Flox/Flox^ (Wnt4^tm1.1Bhr^/BhrEiJ, IMSR_JAX:007032) (Kobayashi et al. 2011)or with Wnt4^GCE/Flox^. Timed matings were checked daily with E0.5 considered as noon of the day the seminal plug was identified. Embryonic kidneys were collected at the indicated times and dissected.

### Immunofluorescence and imaging

For immunohistochemistry of sections, embryonic kidneys were harvested at different time points, fixed overnight in 4% paraformaldehyde at 4°C, paraffin embedded and sectioned in the transverse plane at 6 μm. To minimise inter-slide staining variation, tissue arrays were made by putting 4 to 6 sections from each kidney of two to four embryos on a single slide. Antigen retrieval for all antibodies was carried out using TE buffer (Moreau et al. 2014; Moreau et al. 2019). The antibodies used are listed in Table 1. Finally, slides were mounted in Mowiol with DABCO. Secondary antibodies used for the detection of the primaries are also listed in Table 1. For the TUNEL assay, paraffin sections were stained with Click-iT™ TUNEL Alexa Fluor™ 647 Imaging Assay (Molecular Probes, Thermo Fisher Scientific) according to the manufacturer’s manual.

For wholemount immunohistochemistry, embryonic kidneys were fixed 10-20 mins depending on the developmental stage in PFA 4% in PBS1X. Kidneys were then transferred to a block/permeabilization solution (5% Donkey serum, 0,1% Triton X-100 in PBS 1X). Antibodies were incubated 1-4 days depending on embryonic age. After 1 day of multiple washes in PBS1X with 0.1% Triton X100, samples were incubated with secondary antibodies in PBS1X 0.1% Triton for 1-2 days. After some washes, samples were dehydrated in series of Methanol and cleared using BABB (a mixture of 1 part benzyl alcohol to 2 parts benzyl benzoate; Sigma-Aldrich, benzyl alcohol, cat. no. 305197; benzyl benzoate, cat. no. W213802) (Combes et al., 2014; Moreau et al., 2019). Images were captured on a LSM 980 with airyscan confocal microscope (Carl Zeiss) or a Stellaris 5 confocal microscope (Leica) or two Spinning disk Confocals (3i and DragonFly from Andor).

### Image analysis

Quantification of SIX2 positive cells, LHX1 positive cells and total number of cells on paraffin sections was performed is FIJI (Image J) using StarDist pluggin (Schmidt et al. 2018). The same pluggin was also used to quantify TUNEL and KI-67 positive cells. For HCR, The Fiji macro and CellProfiler pipeline (DOI 10.26180/23628186) were used to analyse the images.

The images were initially processed in Fiji with a customed macro to batch open the lif files (Image taken on Leica Stellaris 5) and split the images into individual channels. The images carried a substantial amount of auto fluorescent structures outside the regions of interest. To eliminate those, the housekeeping gene channel (HKG-C1, *Rnu6*) was thresholded with a Maximum Entropy filter, converted into a binary mask, the Look Up Table (LUT) was inverted to create a mask which was subsequently subtracted from all 5 channels. The resulting images were saved into the user-specified output directory and used as the input for the next processing step in CellProfiler.

The first step of the pipeline was the segmentation of the structures of interest delineated by the housekeeping gene. This was performed by applying the *Identify Primary Object* module (*Thresholding Method > Otsu*; *Typical diameter of objects*> *50-500 pixels*) to the housekeeping image. The output image was named Cells.

The *EnhanceOrSuppressFeatures* module was applied to the 4 channels to enhance the punctated labelling characteristic of RNAscope labelling and eliminate the background noise. The module parameters were set as O*peration > Enhance; Feature type > Speckles; Feature Size > 10 pixels*.

Each resulting images was masked with the Cells objects image to limit the segmentation to the structures of interest labelled with the housekeeping gene. The masked images were further processed with the *Identify Primary Object* (*Thresholding method > Robust Background*). The *RelateObject* module was then used to create a relationship between the Cells objects (*parent*) and the puncta for each gene of interest. This step provides the number of puncta per structure of interest.

The *MeasureOobjectSizeAndShape* modules was applied to all the segmented objects (Cells and genes of interest), the *MeasureImageAreaOccupied* module was applied to the Cells objects. All measurements were exported in an excel spreadsheet.

### Bulk RNA seq and analysis

Total RNA was extracted from freshly dissected embryonic kidneys using the RNAqueous^®^-Micro Kit (Thermo Fisher Scientific). The sequencing was performed at MHTP Medical Genomics at Monash University using the 3’-Multiplex RNA-seq method. Libraries were run on Illumina NextSeq 2000. The resulting count data were imported into R programming language to assemble the gene expression matrix. “biomaRt” R package v.2.46.3 was used to map Ensembl gene Ids to gene symbols. Data were normalized using the CPM method and DEGs were recognized using voom estimated weights and limma R packages and Degust software (BH adjusted p-value < 0.05 & absolute LogFC > 0.5).

### Single cell RNA seq and analysis

Wnt4^GCE/+^ animals were crossed together and Wnt4^Flox/Flox^ were crossed with Six2^TGC^, ^Wnt4Flox/+.^ 23 embryos from 4 females (2 of each cross) were harvested at E14.5. Embryos were sexed and embryonic kidneys were dissected in cold PBS 1X and kept on ice while the samples were genotyped. A piece of embryonic tail was taken from each embryo and DNA was extracted using QuickExtract™ DNA Extraction Solution (Lucigen) according to the manufacturer’s instruction. 1.5 ul of DNA was used in MyTaq™ Red Mix (Bioline) PCR amplification and samples were genotyped using primers from Kobayashi et al. 2008 and 2011. A fast PCR program was developed for this purpose: (95°C, 10 sec - 60°C, 10 sec - 72°C, 15 sec) x 6, (95°C, 2 sec - 60°C, 2 sec - 72°C, 4 sec) x 27 and a final step of 30 sec at 72°C. Amplified DNA were run on agarose gels at 1.2% for 20 mins.

3 pairs of Six2^TGC^, Wnt4^Flox/+^ and 4 pairs of Six2^TGC^, Wnt4^Flox/Flox^ were chosen, and 3 pairs of Wnt4^GCE/GCE^ and 4 pairs of Wnt4^GCE/+^ were pooled per genotyped. Each sample pools were dissociated in 500 ul Accutase (Stem cell Technologies) for 6-7 mins at 37°C and gently mixed by pipetting up and down every 2 mins. Cells were then pelleted by centrifugation at 400g for 5 mins at 4°C and washed in cold PBS 0.04% BSA. Cells were resuspended in 200ul of the same solution and filtered with Flowmi Cell Strainers (70um, Bel-Art Products). Debris, Doublets and dead (DAPI+) cells were removed by FACS on a BD influx (BioScience) using the 100um nozzle.

### HCR

Paraffin sections were deparaffinised and rehydrated. Multiplexed HCR™ RNA-FISH protocol was performed following manufacturer’s instruction (Molecular Instruments). probes were designed by the company and lot number are: HCR amplifier used are B2-Alexa Fluor^®^ 647, B5-Alexa Fluor^®^ 488, B4-Alexa Fluor^®^ 546, B3-Alexa Fluor^®^ 594, B1-Alexa Fluor^®^ 405.

### Kidney culture

Embryonic kidneys were harvested at E12.5 from Wnt4^GCE/+^ parents. Each kidney pair was divided, set up in either vehicle or with BMP4 (314-BP, R&D Systems) and cultured on a transwell insert in a 6-well plate in DMEM with 10% FCS. BMP4 was added at 100ng/ml in 4 mM HCl and 0.1% bovine serum albumin whereas the vehicle condition was only 4 mM HCl and 0.1% bovine serum albumin at the same volume. Treated and untreated kidneys were cultured for 48h, taking them to a developmental stage approximately equivalent to E14.5.

### Droplet Digital PCR (ddPCR)

Embryonic kidneys were dissociated in 500 ul Accutase (Stem cell Technologies) for 8 mins at 37°C and gently mixed by pipetting up and down every 2 mins. Cells were then pelleted by centrifugation at 400g for 5 mins at 4°C and washed in cold PBS 0.04% BSA. Cells were resuspended in 200ul of the same solution and filtered with Flowmi Cell Strainers (70um, Bel-Art Products). Cells were then incubated for 30 mins with LTL (Lotus Tetragonolobus Lectin Biotin conjugated 1:500), washed and then incubated with a Streptavidine 647 (1:2000).

Debris, Doublets and dead (DAPI+) cells were removed by FACS on a BD influx (BioScience) using the 100um nozzle and SIX2-GFP-positive cells as well as LTL-positive cells were retrieved. Cells were then pelleted, frozen down and kept in −80 degrees.

### Droplet Digital PCR (ddPCR) was performed by the Monash Genome Modification Platform (MGMP)

The gene ribonuclease P/MRP 30 subunit (Rpp30) was selected as the reference detection sequence. The primers and probes for digital PCR were designed using Primer3Plus. The ddPCR probe for the reference sequence (Rpp30) was labelled with hexachlorofluorescein (HEX) at the 5’ end and black hole quencher (ZEN/IowaBlack) at the 3’ end (IDT). The ddPCR probe for the target sequence was labeledwith 6-carboxyfluorescein (FAM) at the 5’ end and black hole quencher (ZEN/IowaBlack) at the 3’ end (IDT).

The ddPCR assays were carried out in a total volume of 20 μL containing 20 ng of DNA template, 10 μL2.ddPCR Master Mix (Bio-Rad, USA), 1 μL of 10 μM primer-F, 1 μL of 10 μM primer-R, and 0.5 μLof 10 μM probes. A Bio-Rad QX200 ddPCR droplet generator (Bio-Rad, USA) was used to divide the 20 μL mixture into approximately 20,000 droplets, with the target DNA segments and PCR reagents being randomly distributed between the droplets. The primer and probe sequences are shown in Table 2. The thermal parameters were as follows: 10 min at 95°C to activate the enzyme, followed by 40 cycles of 30 s at 94°C and 1 min at 60°C, followed by enzyme inactivation at 98°C for 10 min and holding at 4°C. The amplified products were analysed using a QX200 droplet reader (Bio-Rad, USA). After the PCR assays were finished, the copy numbers were analysed using QuantaSoftAnalysis Pro Version 1.0 (Bio-Rad, USA).

## Data Availability

Sequencing data presented in this manuscript have been deposited in GEO under accession numbers GSEXXXXXX (bulk RNAseq data) and GSEXXXXX (scRNAseq data).

## Acknowledgements

We express thanks to Rachel Lam, Dr. Kynan Lawlor and Prof. Melissa Little for their support and early contributions to the project. This project was supported by grants from the National Health and Medical Research Council of Australia (APP1156567 to A.N.C.) and the Australian Research Council (DP190101037 to A.N.C.). This project was enabled by equipment and expertise at Monash Micro Imaging, Micromon Genomics, the Monash Histology Platform, Monash Animal Research Platform and Monash Genome Modification Platform (MGMP). We acknowledge the support of Phenomics Australia for MGMP equipment and services. Additional microscopy was performed at the Murdoch Children’s Research Institute. We thank Dr Andrew Perry and the Monash Bioinformatics Platform for assistance, including setup and maintenance of analysis infrastructure on the Nectar Research Cloud. The Nectar Research Cloud is a collaborative Australian research platform supported by the NCRIS-funded Australian Research Data Commons (ARDC). AJM is supported by a Queensland Health Advancing Clinical Research Fellowship.

## Author Contributions

J.L.M. designed and performed experiments, data analysis, presentation, interpretation, writing; S.W. analysis of bulk and single cell RNAseq, writing (methods). J.H-L. image analysis; A.J.M. designed & interpretation, writing; A.N.C. project design, supervision, data analysis & interpretation, writing.

---

**Figure S1:**
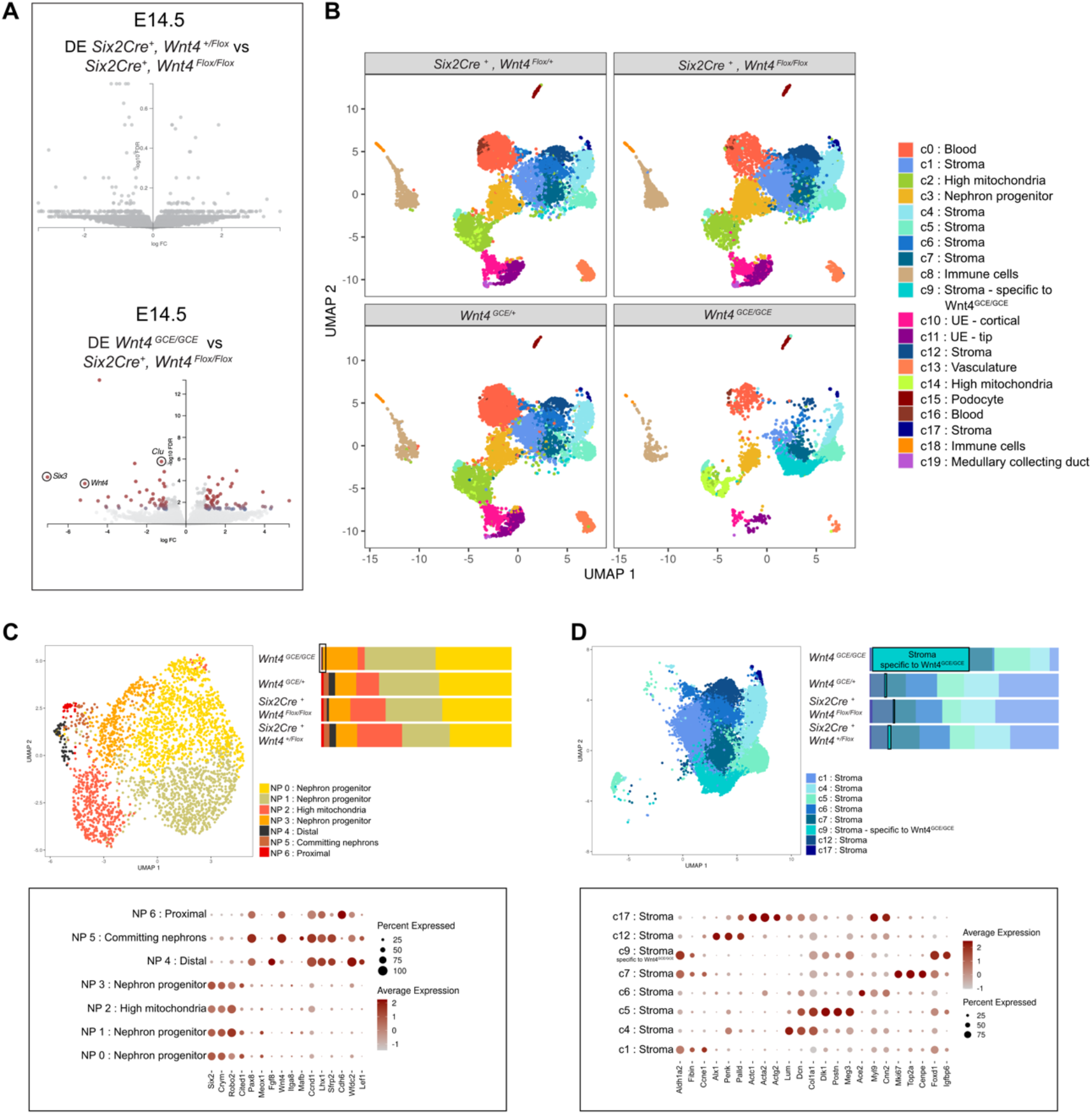
Six2-derived structures and stromal cell populations are transcriptionally affected in absence of *Wnt4*. A) Differential expression volcano plots of Bulk RNA sequencing performed on Six2Cre^+^, Wnt4^Flox/+^,vs Six2Cre^+^, Wnt4^Flox/Flox^ and Six2Cre^+^, Wnt4^Flox/+^ vs Wnt4^GCE/GCE^ at E14.5 B) Separated UMAP of Single Cell RNA sequencing performed on Wnt4^GCE/+^, Wnt4^GCE/GCE^, Six2Cre^+^ Wnt4^Flox/+^ and Six2Cre^+^ Wnt4^Flox/Flox^ kidneys at E14.5 showing the proportion of each cluster per genotype. C) Sub clustering of Nephron progenitors cluster and the relevant markers D) Sub clustering of stromal cell population cluster and the appropriate markers.

